# Automated analysis of sleep in adult *C. elegans* with closed-loop assessment of state-dependent neural activity

**DOI:** 10.1101/791764

**Authors:** Daniel E. Lawler, Yee Lian Chew, Josh D. Hawk, Ahmad Aljobeh, William R. Schafer, Dirk R. Albrecht

**Affiliations:** Department of Biomedical Engineering, Worcester Polytechnic Institute, 100 Institute Road, Worcester, MA 01609 USA; Illawarra Health and Medical Research Institute and School of Chemistry and Molecular Bioscience, University of Wollongong, Wollongong 2522 Australia; Department of Neuroscience, Yale University School of Medicine, P.O. Box 9812, New Haven, CT, 06536-0812, USA; Neurobiology Division, Medical Research Council (MRC) Laboratory of Molecular Biology, Francis Crick Avenue, Cambridge CB2 OQH, UK; Department of Biology and Biotechnology, Worcester Polytechnic Institute, 100 Institute Road, Worcester, MA 01609 USA

## Abstract

Sleep, a state of quiescence associated with growth and restorative processes, is conserved across species. Invertebrates including the nematode *Caenorhabditis elegans* exhibit sleep-like states during development and periods of satiety and stress. Here we describe two methods to study behavior and associated neural activity during sleep and awake states in adult *C. elegans*. A large microfluidic device facilitates population-wide assessment of long-term sleep behavior over 12 h, including effects of fluid flow, oxygen, feeding, odors, and genetic perturbations. Smaller devices allow simultaneous recording of sleep behavior and neuronal activity, and a closed-loop sleep detection system delivers chemical stimuli to individual animals to assess sleep-dependent changes to neural responses. Sleep increased the arousal threshold to aversive chemical stimulation, yet sensory neuron (ASH) and first-layer interneuron (AIB) responses were unchanged. This localizes adult sleep-dependent neuromodulation within interneurons presynaptic to the AVA premotor interneurons, rather than afferent sensory circuits.

## Introduction

Sleep is a physiological state during which voluntary muscle activity ceases, sensory processing is modulated^1^, and anabolic, growth, and restorative processes occur in the brain and other tissues^2^. Sleep is observed across species, from mammals to invertebrates^3^, where it controls energy usage^4^, metabolism, macromolecular biosynthesis^5^, and neural plasticity and memory consolidation^6^. Sleep pathologies include improper duration or control (e.g., insomnia, narcolepsy), altered sleep behavior (e.g., sleepwalking), and altered sensation (e.g., restless leg syndrome). In humans, these sleep deficiencies are associated with reduced productivity^7^ and increased prevalence of cardiovascular disease^8^, diabetes^9^, and obesity^10^.

The initiation and cessation of sleep is often mediated by circadian rhythms, which are controlled by environmental factors^11^ and timing genes that are generally conserved across species^12^. A number of molecular pathways^13–19^ are involved in promoting sleep states and inhibiting arousal behavior. However, it is currently unclear how these pathways modulate circuit-level sensory processing during sleep states^20^, and how misregulation of neural activity may contribute to sleep disorders.

The nematode *C. elegans* provides distinct advantages for direct observation of neurological function in freely-behaving animals. They are small (<1 mm), exhibit short generational times, and have a compact and fully mapped connectome of 302 neurons in hermaphrodites and 387 in males^21^. Noninvasive optical measurements of neural activity can be made in living, behaving animals via genetically-encoded fluorescent calcium indicators such as GCaMP^22^, and genetic tools are available for rapid generation of mutants and transgenic strains for mechanistic studies^23–25^.

*C. elegans* demonstrate states of quiescence during lethargus between larval stages^26^ (developmental sleep) and during periods of stress^27^, satiety^28,29^, starvation^30,31^, and hypoxia^32^. Additionally, adult *C. elegans* undergo quiescent periods after 1–2 h of swimming in liquid^33^ and in microfluidic chambers with open and constrictive geometries^34^. These quiescent states share fundamental characteristics with sleep in other species^35^, including putative human sleep functions such as processing of synaptic plasticity^36^ and metabolic control^37^, and typical behavioral characteristics such as increased arousal threshold^26,38^, stereotypical posture^39–41^, homeostatic response to sleep deprivation^26,37,42^, and rapid reversibility^26,43^. Together, these connections make a strong case that these quiescent bouts represent sleep in *C. elegans*^44^, enabling its use as a model organism to study the molecular basis of sleep.

*C. elegans* sleep has been observed on a variety of experimental platforms, including agar^26^ or agarose pads^45,46^ and microfluidic chambers that house individual animals^34,47–49^ throughout multiple development stages. Agar pads are useful for studying developmentally-timed sleep, by permitting feeding and growth across multiple larval stages, but they are difficult to use for assessing neural responses to sensory stimulation. Neural activity measurements typically require immobilization by agarose pads^50^ or microfluidic traps^38^, which prevent locomotory behavior and limit their use to developmentally-timed sleep in which sleep state is inferred by timing rather than by behavior. Alternatively, whole brain imaging in immobilized adult animals^31,32^ showed cessation of spontaneous neural activity during presumed sleep states induced by hypoxia. However, spontaneous sleep events in adult animals are best assessed by analysis of locomotion and quiescent behaviors. Therefore, new methods are needed for monitoring sleep state and stimulated neural responses in freely-moving animals, in order to assess the functional circuit changes that occur during adult sleep.

Here we demonstrate two systems to quantify the behavioral and neural characteristics of sleep in young adult *C. elegans*, using microfluidic devices that allow a natural sinusoidal crawling motion in a well-controlled liquid environment enabling rapid chemical stimulation^51^. A larger device is used for population-wide assessment of over 100 animals and up to 4 conditions at once, and a smaller device allows simultaneous behavioral and neural recording in individual animals^52^. We show that sleeping behavior exhibited by young adult *C. elegans* follows characteristic dynamics over 12 h in microfluidic devices and is impacted by fluid flow, oxygen, bacterial food, food signals, and genetic perturbations affecting sensory input. In this platform, the onset of adult sleep correlates with increased activity of the RIS interneuron^53,54^. Using a closed-loop chemical stimulation system, we observed an increased arousal threshold during adult sleep states as before^26^ and also monitored simultaneous neural activity. A sleep-dependent delay in response to aversive copper chloride corresponded to diminished and delayed responses in AVA command interneurons. Responses in the ASH sensory neurons and AIB interneurons were not modulated by sleep in young adult animals, localizing sleep-state neural circuit modulation within interneurons of the aversive sensorimotor subcircuit. These results suggest that sleep specifically alters the linkage between sensory stimuli and command neurons without changing processing within high-level interneurons, and provide an experimental system to dissect the molecular processes that produce this specificity.

## Results

### High-throughput analysis of adult sleep

Sleep behavior, defined by periods of behavioral quiescence, was observed in young adult *C. elegans* over 12 hours in microfluidic behavior arenas^51^. A hexagonal array of 70 μm tall microposts enables free sinusoidal crawling behavior in a dynamic, switchable liquid environment with continuous flow (**Fig. 1a,b**). Each device contained four 16 mm × 15 mm arenas housing four independent populations of ~25 freely crawling animals that share the same fluidic environment. Wild-type animals roam around microfluidic arenas with predominantly forward locomotion, separated by momentary pauses, spontaneous short reversals (<1 s), and long reversals coupled with reorienting “omega” turns^51,55^. Awake animals may pause briefly to feed if bacterial food is present^56^, to probe the physical barriers of arenas, or when encountering other animals. Other times, animals enter a prolonged quiescence state that lasts for ~20 s to several minutes (**Supplementary Video 1**). These bouts begin with animals gradually slowing their mean forward locomotion speed over 10–20 s (**Fig. S1a**), often pausing briefly a few times (**Fig. S1b**) during slow, creeping motion. Animals then gradually adopt a relaxed body posture^41^ over about one minute (**Fig. S1c,d**) and cease further movement. Sleeping animals are apparent visually in microfluidic arenas by their straight head and contact with only one micropost (**Fig. S1d**), whereas awake animals actively wrap around several posts (e.g., **Fig. 1b**). After typically one or more minutes, animals quickly wake and resume forward (or occasionally reverse) locomotion, accelerating to a typical 0.15 mm/s forward velocity within 5 s.

**Figure 1.**
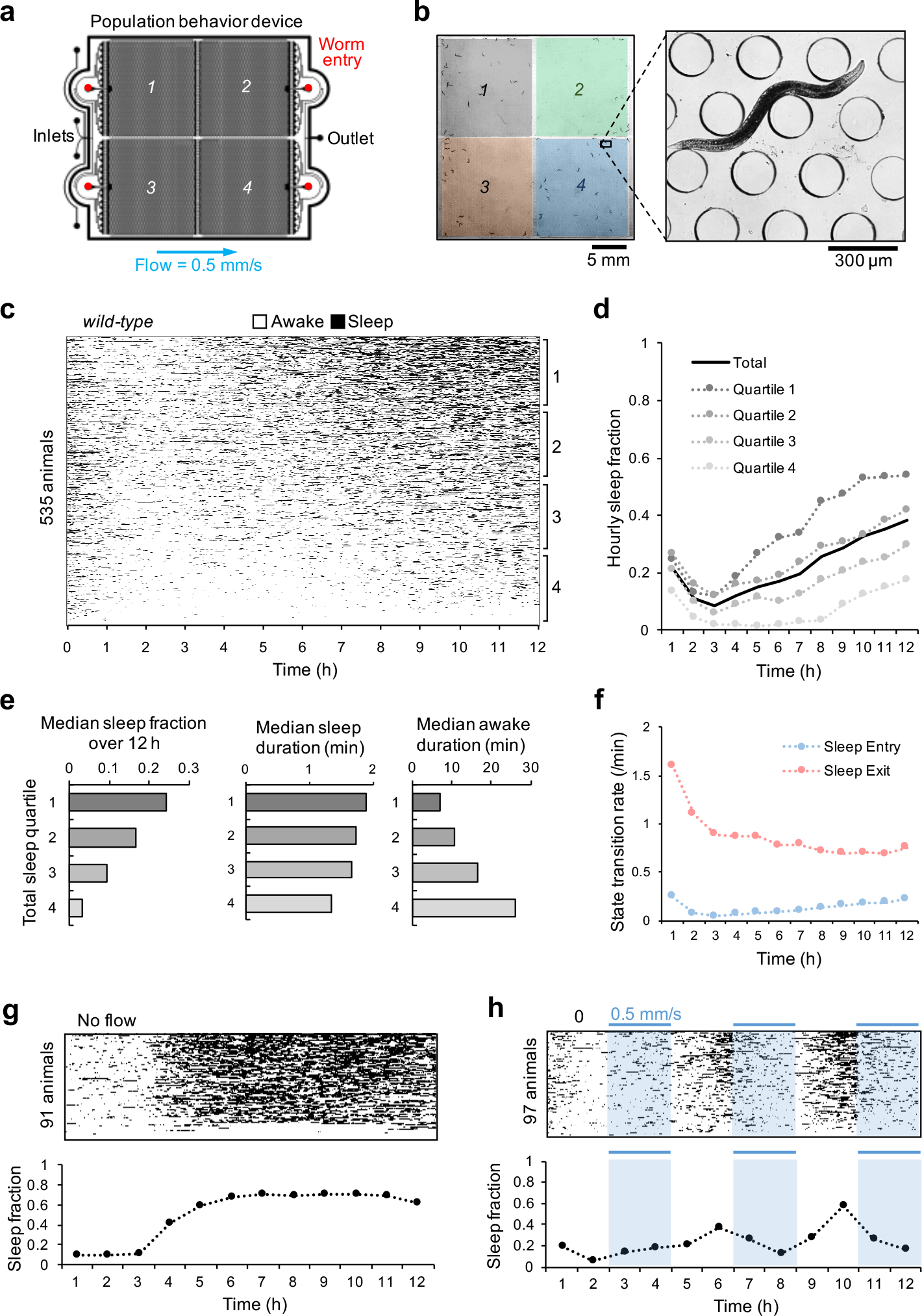
Young adult sleep in wild-type *C. elegans* in “Population behavior” microfluidic devices. (**a**) Schematic of the “Population behavior” microfluidic device, including multiple inlets to switch fluids, four worm entry ports to introduce separate worm populations, and a flow outlet. (**b**) Image frame of a device containing about 100 animals, 25 in each of four separated 16 mm × 15 mm arenas. Awake animals roam freely between 200 μm diameter microposts (inset). (**c**) Heatmap of sleep events (black) over 12 h, sorted by total sleep fraction (*n* = 535 animals). (**d**) Hourly sleep fraction for all animals from **c** and grouped into four quartiles by their total 12 h sleep fraction as in **c** (quartile 1 = most sleep). (**e**) Median sleep fraction, sleep bout duration, and awake bout duration from data in **c**, separated by total sleep quartiles as in **c**. (**f**) Sleep entry/exit transition rate plotted during each hour of experimentation from data in **c**. (**g**) Effect of stationary fluid on sleep behavior over 12 h (*n* = 91 animals). Heatmap of sleep events shown above, and mean hourly sleep fraction below. (**h**) Effect of pulsed buffer flow on sleep behavior over 12 h (*n* = 97 animals). Fluid flow alternated between moderate flow (0.5 mm/s) or no flow every 2 h. Heatmap of sleep events shown above, and mean hourly sleep fraction below.

Since pauses reflect both the extended quiescent states of sleep bouts and the momentary pauses of awake animals, true sleep states were automatically identified by centroid tracking filtered by the characteristic duration, history, and body shape of sleep. Using temporal parameters based on sleep entry and exit dynamics (onset after 20 s continuous pausing and ending at 5 s non-pausing), automatic classification of sleep bouts showed 95.2% agreement with human observation, with slight underestimation of sleep states (1.9% false discovery rate; 7.9% false omission rate; *n* = 500 randomly selected bouts; **Fig. S1e**). These detected sleep bouts exclude the brief pauses that precede a sleep bout, and include momentary “twitch” movements during sleep which can be caused by contact from other animals, flow disturbance, or presumed involuntary movements, and do not signal exit of a sleep state.

We analyzed 535 wild-type (N2) animals for 12 h in continuous 0.5 mm/s flow of S. Basal buffer (**Fig. 1c**). Hourly sleep fraction, defined by the fraction of time the animal spends in a sleep state during each hour, decreased on average across the population from 22% ± 0.8% s.e.m. in the first hour to 8% ± 0.5% in hour 3, then increased steadily to 38% ± 1% in hour 12 (**Fig. 1d**). A wide range of sleep behavior was observed among individual wild-type animals, with 95% exhibiting a 12 h total sleep fraction ranging from 4% to 43%. To assess variability in sleep dynamics, we divided animals into quartiles by total sleep fraction. Sleep dynamics were similar in all quartiles, with sleep fraction increasing over time after 3 h (**Fig. 1d**), but median sleep fraction over 12 h varied greatly across quartiles from 3% to 24%. Median sleep duration remained between 1.3–1.9 min for each quartile (**Fig. 1e**), whereas median awake duration varied more greatly, with the top quartile of sleeping animals remaining awake for a median of 7 min, about one-quarter of the most active animals (27 min). The increase in sleep fraction over time from hours 3–12 was associated with both an increased rate of sleep entry (more sleep pressure) and a decreased rate of sleep exit (more sleepiness) (**Fig. 1f**), resulting in more frequent and longer sleep bouts and shorter awake periods over time (**Fig. S2a**). The rate of sleep exit remained consistent across total sleep quartiles (**Fig. S2b**), while the rate of sleep entry varied greatly (**Fig. S2c**). Together, these results indicate that sleep bouts were similar across wild-type animals, while individual variability in sleep fraction among wildtype animals predominantly arose due to variation in the rate of sleep entry, or equivalently, the duration of awake bouts, and in the frequency of sleep bouts.

### Environmental and sensory effects on sleep dynamics

Sleep entry and exit are sensitive to environmental conditions and sensory input. To test the role of sensory input on sleep, we first assessed the effect of flow in the microfluidic environment, comparing sleep amounts with a moderate flow rate (0.5 mm/s), no flow, and periodic pulsing of flow conditions (**Fig. 1g,h**). Without flow, sleep fraction was similar to moderate flow conditions for the first 3 h, but rose dramatically from 12% to 42% around 3 h and remained high for the duration of the 12 h experiment (**Fig. 1g**). To test whether resumption of flow would return sleep fraction to baseline rates, we pulsed flow every 2 h, alternating between 0.5 mm/s flow or no flow. Again, sleep fraction remained low for the first 3 h regardless of flow condition, then sleep fraction became flow-dependent, increasing after about 30 mins without flow and decreasing sharply when flow was resumed (**Fig. 1h**). Under static conditions, animals can deplete the microfluidic environment of oxygen^57,58^, and hypoxia has been shown to induce sleep behavior^59^ especially in starved animals^31^.

We therefore assessed the role of oxygen in adult sleep in microfluidic devices. With continuous flow of 0.5 mm/s, a hypoxic buffer (<1% O_2_, 30 mM sodium sulfite^60^) significantly increased total sleep fraction (48% ± 1.1%, P<0.0001) compared to the same solution reoxygenated to >20% O_2_ (16% ± 1.3%) (**Fig. 2a-c**). During hypoxia, 13% of sleep bouts were >10 min long, including some bouts lasting hours, compared to only 3.5% in the reoxygenated buffer (**Fig. S2d**). Notably, hypoxia increased sleep fraction only after 4 h in the device (P<0.0001, **Fig. 2c**), in line with past results suggesting that starvation and hypoxia work together to promote sleep behavior^31^. The rapid rise in sleep behavior after 4 h mimicked a similar rise in static no-flow conditions (**Fig. 1g**), suggesting that gentle flow replenishes oxygen to suppress sleep behavior.

**Figure 2.**
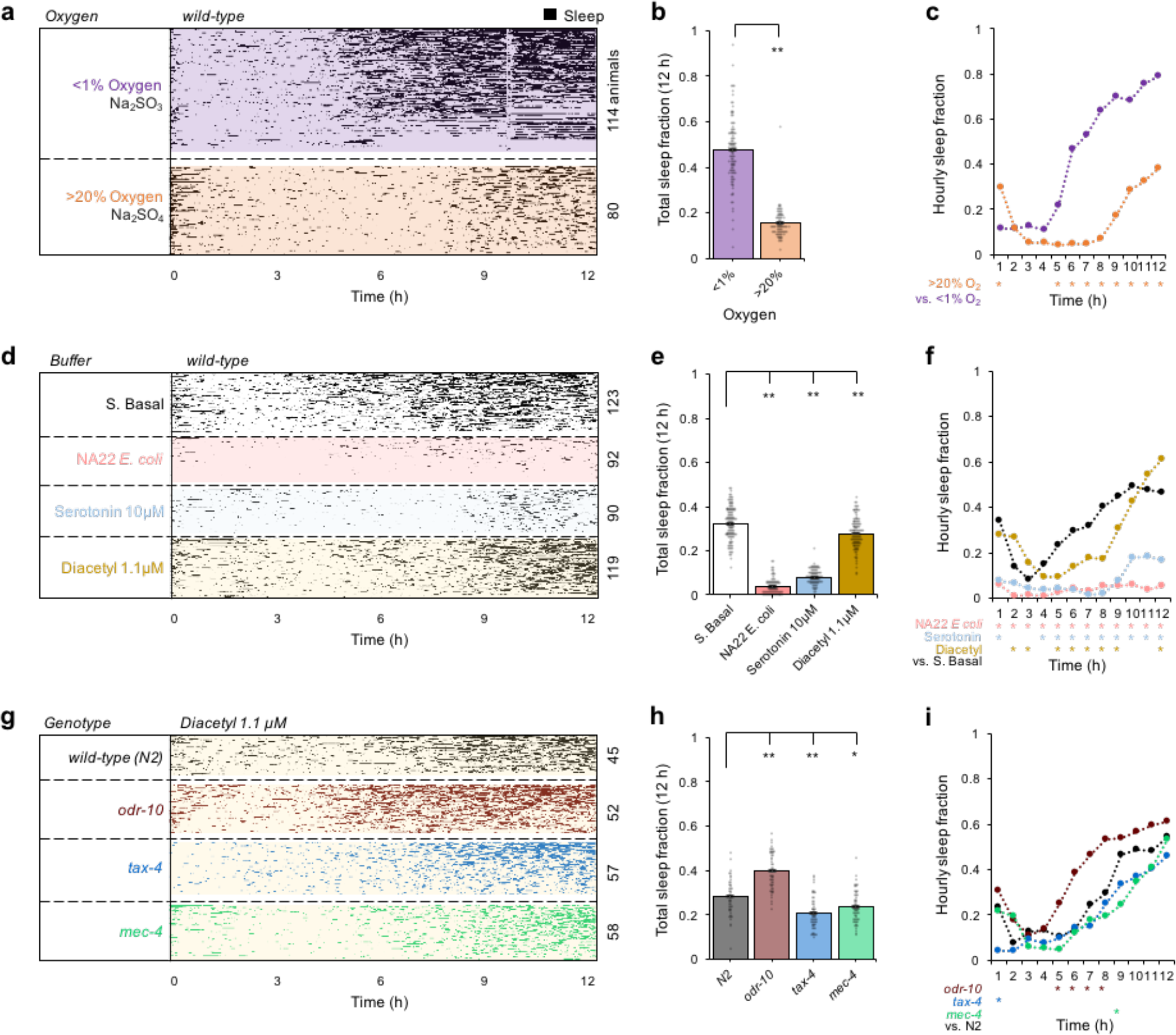
Effects of oxygen, feeding, food signals, and sensory input on young adult sleep dynamics. (**a**) Effect of hypoxia on sleep dynamics (*n* = 80–114 animals). Within each group, animals (heatmap rows) are sorted by total sleep fraction. (**b**) 12 h total sleep fraction assessing effect of hypoxia on sleep behavior from **a** with bars representing population mean ± s.e.m. and points indicating individual animals. (**c**) Hourly sleep fraction from data in **a**. (**d**) Effect of feeding and food signals comparing sleep behavior in S. Basal buffer vs. bacterial food (NA22 *E. coli*, OD600 = 0.35), serotonin 10 μM to mimic feeding response, and a food odor diacetyl 1.1 μM (*n* = 90–123 animals). (**e**) 12 h total sleep fraction assessing feeding effect on sleep behavior, as in panel **b**. (**f**) Hourly sleep fraction from data in **d**. (**g**) Sensory mutant sleep behavior assessed in diacetyl 1.1 μM (*n* = 45–58 animals). (**h**) 12 h total sleep fraction assessing effect of sensory mutations on sleep behavior, as in panel **b**. (**i**) Hourly sleep fraction from data in **g**. Statistics for all plots were performed using one-way ANOVA with Bonferroni’s correction for multiple comparisons. For 12 h total sleep fraction plots (b, e, h): ** P<0.0001; * P<0.05. For hourly sleep fraction (c, f, i), significance is noted for * P<0.0001 as indicated within data of that hour.

Because feeding state impacts arousal^61,62^ and starvation may regulate the impact of hypoxia on sleep^31^, we next assessed the role of feeding and satiety on adult sleep dynamics within our microfluidic chamber (**Fig. 2d-f**). The presence of bacterial food (NA22 *E. coli*) suppressed total 12 h average sleep fraction (3.8% ± 0.6%, P<0.0001) compared to S. Basal control (33% ± 0.5%) (**Fig. 2e**). Serotonin, which mimics the feeding response^63^, presented at a moderate concentration of 10 μM similarly reduced total sleep fraction (8% ± 1.1%, P<0.0001) compared to control buffer conditions. Whereas bacterial food suppressed sleep continuously for 12 h, serotonin suppressed sleep for ~9 h. Similarly, a moderate behaviorally attractive food odor^64^ (1.1 μM diacetyl) suppressed total sleep fraction compared to control buffer (28% ± 0.9%, P<0.0001), although to a lesser extent than food or serotonin. Diacetyl suppressed sleep fraction up to hour 9 (P<0.0001), consistent with adaptation to the odor over hours^65,66^ (**Fig. 2f**). Animals also showed increased hourly sleep fraction when starved on a plate without food prior to entry to the microfluidic environment (**Fig. S3a**). These results suggest that adult sleep behavior in microfluidic devices is driven in part by feeding state and perception of hunger.

To observe how sensory information influences sleep, we tested wild-type animals and three sensory mutants (**Fig. 2g-i**) loaded into separate arenas of each “Population behavior” device (**Fig. 1a**). Since the odorant diacetyl reduced sleep (**Fig. 2f**), we tested *odr-10* mutants, which lack the diacetyl receptor normally present in the AWA sensory neurons and should not perceive this odor. In the presence of 1.1 μM diacetyl, *odr-10* mutants exhibited a higher total sleep fraction (40% ± 1.0%, p<0.0001) compared to wild-type (28% ± 1.2%) (**Fig. 2h**), and similar to wild-type animals in control conditions (**Fig. S3b**). In diacetyl, *odr-10* mutants showed a significant increase in hourly sleep fraction compared to wild-type only up to 6 h (**Fig. S3c**), after which habituation to the odor may reduce its influence. Sensory deficient *tax-4* mutants lack a cyclic GMP-gated ion channel necessary for signal transduction in many sensory neurons^67^ and are defective in multiple sensory behaviors, failing to respond to temperature or to water-soluble or volatile chemical cues. However, *tax-4* is not present in AWA neurons; hence, diacetyl-mediated sleep suppression should be preserved in this mutant. Indeed, while *tax-4* showed a moderate decrease in total sleep fraction (20.7% ± 1.0%, P<0.0001) compared to wild-type over 12 h in 1.1 μM diacetyl, no significant differences in hourly sleep fraction were observed except during the first hour (**Fig. 2i**). Strong suppression of early quiescence bouts in hour 1 in *tax-4* animals (4% vs. 21%, **Fig. S3d**) suggests that sensory information other than from AWA neurons contributes to elevated quiescence in the first hour. Animals transferred into microfluidic devices experience a novel mechanical environment, including gentle touch of the microposts and continuous fluid flow. While gentle touch deficient *mec-4* mutants showed a slightly lower total sleep fraction than wild-type (24% ± 1.0% vs. 28% ± 1.2%, P<0.05), *mec-4* mutants had no significant difference in first hour sleep fraction compared with wild-type (18% vs. 21%, **Fig. S3d**), suggesting that any sensory information leading to elevated initial quiescence did not come from the *mec-4*-expressing touch receptor neurons ALM, AVM, or PLM. Together, these data demonstrate the role of sensory information in sleep regulation, and the testing of multiple mutants at once in multi-arena microfluidic devices to investigate regulators of sleep dynamics.

### Automatic sleep tracking, chemical stimulation, and neural imaging

To understand how neural activity changes during sleep cycles, we designed smaller “Neural imaging” microfluidic devices, containing a single 3 mm × 3 mm arena with the same micropost array as the “Population behavior” device (**Fig. 3a**) but sized to fit the entire field of view at 5× magnification on an epifluorescence microscope (**Fig. 3b**). Sleep behavior of animals was tracked using brightfield illumination every 10 s, using frame subtraction similar to previous methods^48^ (**Fig. S4a,b**), and correctly identified sleep bouts with 93.4% agreement with human observers (**Fig. S4c**). Wild-type *C. elegans* sleep dynamics in the small “Neural imaging” device were equivalent to the larger “Population behavior” devices, dropping from 18% to 6% over the first 3 h, then steadily rising to 50% by 12 h, despite a faster flow velocity in the neural imaging microfluidic device (15 mm/s vs. 0.5 mm/s) (**Fig. 3c**).

**Figure 3.**
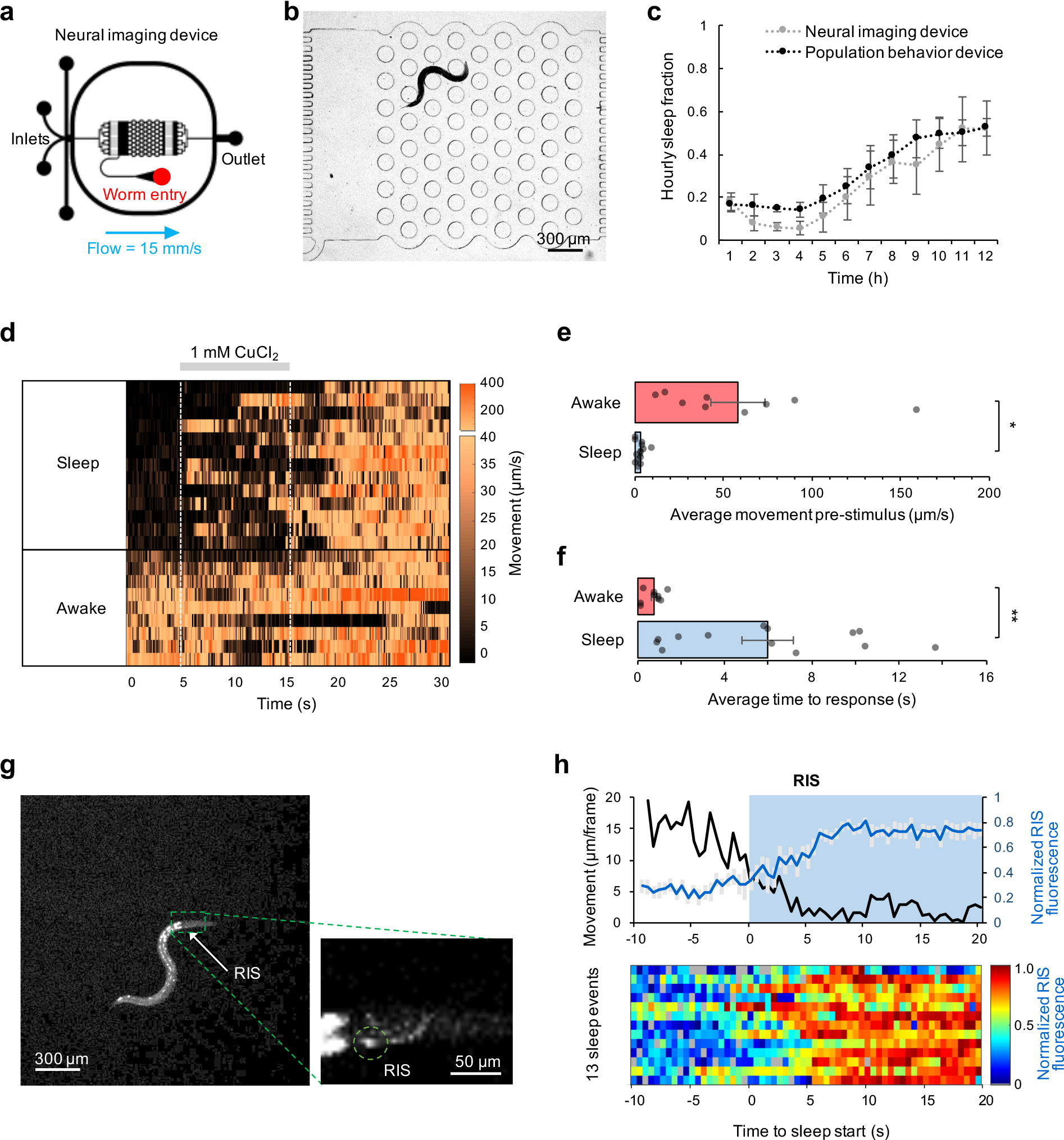
Neural activity during sleep and awake states in “Neural imaging” microfluidic devices. (**a**) Design of microfluidic device for closed-loop sleep assessment, chemical stimulation, and neural imaging. Device contains a single 3 mm x 3 mm arena. (**b**) Individual animals tracked using pulsed brightfield illumination (λ = 520–550 nm). An awake animal is shown. (**c**) Comparison of sleep fraction in “Neural imaging” (*n* = 7 animals) vs. “Population behavior” (*n* = ~100 animals) devices. (**d**) Heatmap showing movement of AIB neuron per frame (0.1 s) across 22 pulsed stimulation trials (rows). Stimulus of 1 mM CuCl_2_ applied between 5–15 s during each 30 s trial. Data sorted by average movement 5 s prior to stimulus, indicating the sleep/awake state for each recording. (**e**) Average movement pre-stimulus (0–5 s) in **d** grouped by sleep or awake state (*n* = 13 Sleep, *n* = 9 Awake). (**f**) Average time to a reversal or avoidance behavior response in **d** for sleeping and awake animals. Statistics for **e** and **f** performed using an unpaired two-tailed t-test; ** P<0.001; * P<0.05. (**g**) Image of an animal expressing GCaMP in the RIS neuron during sleep onset. (**h**) Average RIS neuron fluorescence (*n* = 13 traces) and average neuron centroid movement per frame (2 fps). Neural activity normalized to minimum and maximum intensity of each RIS neuron trace during the 30 s before and after the awake to sleep transition at time = 0 s. Heatmap of all neural recordings shown below.

The “Neural imaging” device provides fast temporal control of chemical stimuli, capable of reproducible fluid switching in <0.5 s (**Fig. S4d**) without disturbing natural behaviors. We assessed arousal from sleep by testing sensory responsiveness of sleeping and awake wild-type animals to aversive 10-s pulses of 1 mM copper chloride solution. Sleep or wake states were determined by average pixel movement 5 s prior to stimulation (**Fig. 3d**), which was significantly higher in awake vs. sleeping states (5.8 ± 1.5 μm/frame vs. 0.32 ± 0.07 μm/frame, P<0.05) (**Fig. 3e**). We recorded the time elapsed between chemical onset and the first reversal movement. Responses in a sleep state were about eight times slower (6.0 s ± 1.2 s, P<0.001) than in an awake state (0.76 s ± 0.14 s), consistent with an increased threshold for sensory responsiveness in sleeping young adult animals (**Fig. 3f**) as has been shown during lethargus to mechanical and chemical stimuli^26^.

RIS interneuron activity correlates with the onset of developmentally-timed sleep^53^ and quiescent behavior in adults^54^. To demonstrate neural imaging during spontaneous sleep-wake cycles in the microfluidic device, we recorded activity in the RIS interneuron expressing GCaMP3 (**Fig. 3g**) in freely moving animals while simultaneously assessing movement behavior. As expected, RIS activity increased at the onset of adult sleep (**Fig. 3h**).

### Closed-loop stimulation and neural imaging of a reversal circuit

An increased threshold for sensory responsiveness during sleep suggests sleep-dependent modulation to neural activity in *C. elegans*, either in sensory responses to stimulation, or in downstream interneurons or motor neurons. For example, diminished ASH sensory neuron activity to aversive chemical pulses (1 mM copper chloride) was reported during lethargus states in developmentally-timed sleep^38^. However, it is unclear whether sensory-level modulation occurs during adult sleep as well. Since adult sleep is not synchronized across animals, or within an individual, we developed a closed-loop system that monitors sleep state every 10 s and triggers a stimulation and neural recording when user-programmable conditions are met (**Fig. 4a, Supplementary Video 2**). Here, we chose to stimulate one minute after a sleep state transition, allowing a 15-minute recovery period between stimulation trials (**Fig. 4b**). Brief pulses of blue light excitation were used for fluorescent imaging to measure calcium activity during each 30-s trial (**Fig. 4c**), as strong blue light can cause arousal by itself^68^, and sleep state was monitored by behaviorally-neutral green light.

**Figure 4.**
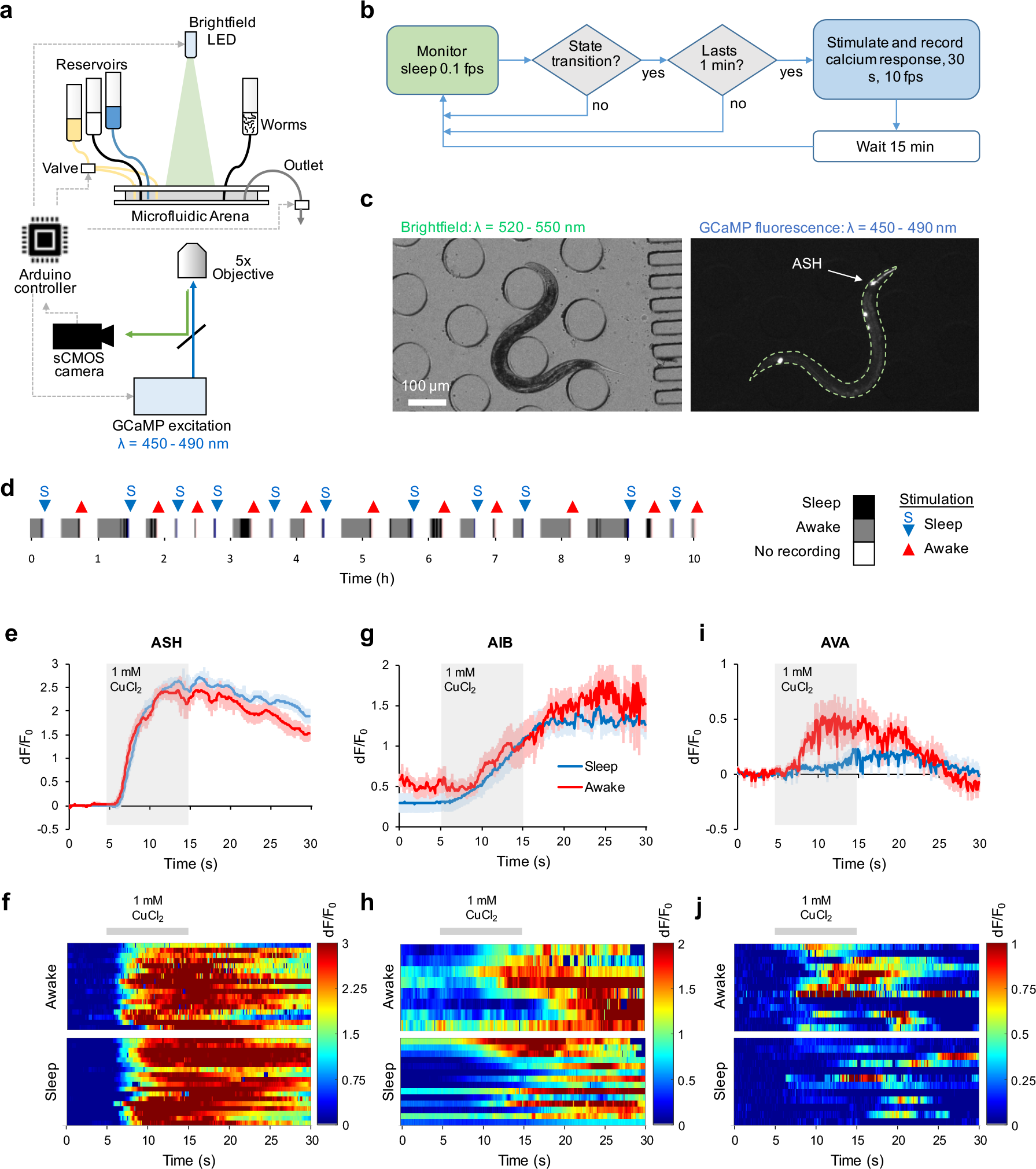
Closed-loop stimulation and neural recording in individual free-behaving animals. (**a**) Schematic of closed-loop neural recording set-up for sleep/awake response tracking. Video recording, valve control, and LED triggering are controlled through an Arduino microcontroller. Brightfield images are used for tracking sleep behavior and fluorescent images are used for measuring calcium transients. Image capture, sleep/awake determination, and chemical stimulation are controlled by computer in a closed-loop without user intervention. (**b**) Decision process schematic of closed-loop experiment. Green (brightfield) and blue (fluorescent) shading of decision nodes indicate corresponding illumination source during frame capture. (**c**) Brightfield (λ = 520–550 nm) and fluorescent (λ = 450–490 nm) images of a freely-moving animal expressing GCaMP in ASH neurons. (**d**) Example showing behavior and neural recording trials in a typical 10 h closed-loop experiment. (**e**) Average ASH neural responses in Sleep and Awake states to 10 s CuCl_2_ pulses (*n* = 18 Sleep, 17 Awake). (**f**) Heatmap of individual ASH responses from **e**. (**g**) Average AIB neural responses in Sleep/Awake states to 10 s CuCl_2_ pulses (*n* = 13 Sleep, 8 Awake). (**h**) Heatmap of individual AIB responses from **g**. (**i**) Average AVA neural responses in Sleep/Awake states to 10 s CuCl_2_ pulses (*n* = 13 Sleep, 12 Awake). (**j**) Heatmap of individual AVA responses from **i**.

We measured neural responses to 10-s pulses of 1 mM copper chloride in the ASH sensory neurons over 12 h in individual animals. A typical closed-loop experiment with 15-minute recovery per stimulation showed no significant adaptation (**Fig. S5a**) and recorded about one sleep and one awake response per hour over >10 hours (**Fig. 4d, Fig. S5b**). ASH neurons responded strongly and consistently to each copper chloride pulse, regardless of sleep or awake state during stimulation (**Fig. 4e,f**).

Since ASH chemosensory responses were equivalent in sleep and awake states, the elevated arousal threshold in sleep could result from diminished activity in interneurons, motor neurons, or in the muscles themselves. ASH is directly presynaptic to AVA, but also has secondary connections through AIB, AVD, and RIC interneurons (**Fig. S5c**). As AIB shares a gap junction with the sleep-inducing neuron RIS, and ablation of AIB reduces long reversals^55^, we recorded AIB and AVA neural activity in sleep and awake states in response to 1 mM copper chloride. Neural responses in AIB were not significantly different between awake and sleep states (**Fig. 4g,h**). In contrast, animals in sleep states had diminished AVA responses, increasing average relative GCaMP fluorescence 55% when awake and 23% when asleep (P<0.05). AVA neural responses were also delayed relative to the copper pulse (**Fig. 4i,j**), consistent with delayed and shortened reversal behaviors (**Fig. 3f**). AIB activity often increased before reversal behavior in sleeping animals but coincided with reversal responses in awake animals (**Fig. S5d**), suggesting a sleep-dependent behavioral delay downstream of (or bypassing) AIB and presynaptic to AVA, that contributes to the apparent arousal threshold increase in sleeping animals.

## Discussion

*C. elegans* sleep has been studied previously during developmentally-timed transitions (lethargus) and after induction by satiety or various stresses. Spontaneous adult sleep has been technically more difficult to assess. Here we present two microfluidic tools to study spontaneous or induced sleep in young adult *C. elegans*, complimentary to related methods for use in larval stages. Both devices share the same microfluidic arena geometry and elicit the same sleep dynamics over 12 h. The larger “Population behavior” device quantifies sleep dynamics to compare up to four different environmental conditions or genetic perturbations at once. The smaller “Neural imaging” device identifies sleep state-dependent neural circuit modulation, by correlating simultaneous recordings of sleep behavior and stimulated neural responses in individual animals. Unlike previous microfluidic methods, these designs used microposts structured for natural crawling behavior without mechanical constriction. Some features of spontaneous adult sleep in this environment differ from previous studies, such as the role of oxygen and flow, and it remains to be seen whether this mode of sleep behavior is unique to stress-induced sleep states previously observed.

Adult quiescence behavior in these devices displays several common behavioral characteristics of sleep. For example, quiescent adults exhibit: (1) an increase in arousal threshold to an aversive chemical stimulus by a delay in behavioral response (**Fig. 3f**), (2) rapid sleep reversibility upon changes in fluid flow (**Fig. 1h**), and (3) a characteristic relaxed posture (**Fig. S1c,d**). Additionally, our results are consistent with (4) a homeostatic sleep response, in which periods of elevated sleep are followed by reduced sleep, and vice versa. For example, sleep suppression by diacetyl odor elicited a higher fraction of rebound sleep at later times (46% ± 1.4% for control buffer alone vs. 57% ± 1.4% for diacetyl in hours 10-12, P<0.0001, **Fig. 2f**, **S3f**). Conversely, elevated sleep during static fluid conditions later suppressed sleep after flow was resumed (21% ± 1.9% for pulsed flow vs. 37% ± 0.8% for continuous flow in hours 10-12, P<0.0001, **Fig. 1h**, **S3f**).

Sleep behavior is sensitive to environmental conditions presented in microfluidic devices. For example, fluid flow in the microfluidic environment is important for maintaining a fresh and constant environment, and cessation of flow increased sleep behavior dramatically. Static fluid conditions may decrease mechanical stimulation, deplete nutrients and oxygen, and increase concentrations of byproducts and CO_2_. As sleep behavior increased dramatically in a hypoxic buffer, oxygen depletion by animals may be a primary factor driving elevated sleep in static microfluidic conditions. While hypoxia increased sleep behavior only after 4 h in freely-behaving animals, sudden hypoxia was shown to suppress most spontaneous neural activity across the whole brain of trapped *C. elegans* in 16 h starved animals^31^. In mammals, intermittent hypoxia may cause excessive sleepiness^69^, but also cause disturbed and superficial sleep with frequent waking via chemoreceptor reflex pathways^70^. Thus, there is an interplay between arousing and somnolent environmental cues. Further studies in *C. elegans* may be useful to distinguish between these contrasting hypoxic effects and to understand the role of sleep in regulating metabolic and energic systems.

Sleep behavior is also dependent on the availability of food. Wild-type animals in buffer without a food source increased their sleep fraction over time away from food, due to both increased “sleepiness” extending sleep bouts and increased “sleep pressure” shortening awake bouts. Feeding bacteria in the device suppressed sleep bouts for at least 12 h, whereas presenting exogenous serotonin to mimic feeding response or a food odor suppressed sleep for 9 h, consistent with adaptation to these food signals.

Sensory neural activity also modulates sleep. For example, sleep suppression by diacetyl was absent in *odr-10* mutants that lack only the diacetyl odor receptor and are unable to detect this odor. Sensory information also contributes to the initial elevated sleep behavior seen in the first hour of testing as animals acclimate to the microfluidic environment. The general sensory mutant *tax-4* suppressed first-hour sleep whereas mechanosensory-deficient *mec-4* animals did not, suggesting that sensation of flow or gentle touch of microfluidic structures do not contribute to early sleep behavior. Instead, other *tax-4* dependent sensation, such as from various thermo- and chemosensory neurons^67^, may be involved in detecting the novel microfluidic environment.

We observed freely-moving behavior and simultaneous activity of several neurons during sleep and awake states. We verified the RIS interneuron is active at the onset of spontaneous adult sleep, as has been shown during developmentally-timed lethargus sleep^53^. The automated closed-loop stimulation system operates without user intervention, resulting in unbiased measurements of neural responses during alternating sleep and awake bouts within the same animal. The ability to record response differences in individuals over time is particularly important given the wide variation in sleep dynamics observed across individual animals. Isogenetic animals, even when raised on the same plate from the same parental animal, exhibited total sleep fractions varying from zero to nearly one half over 12 h. Given the sensitivity of adult sleep to oxygen, feeding state, chemicals, and likely other sensory stimuli, it is possible that even animals cultured identically experience slight variations in their sensitivity to these parameters to induce differences in their sleep dynamics. The ability to automatically monitor individual animals allows for longitudinal studies capturing dozens of events per animal to identify intra-animal differences in sensory processing irrespective of population-wide variation in sleep patterns.

An increased arousal threshold in sleeping animals suggests modulation to sensorimotor neural circuit activity in *C. elegans* during sleep. Responses of the AVA command interneurons, which are required for backward locomotion^55,71,72^ were indeed diminished and delayed during adult sleep, coinciding with delayed behavioral responses. Similarly, diminished AVA activity was previously observed during lethargus^38^. However, sensory responses in ASH neurons were not modulated by sleep state in adults, in contrast to the lower ASH responses observed in larval stages during developmentally-timed sleep^38^, suggesting that spontaneous adult sleep is a distinct phenomenon. The first layer interneuron AIB, which shares synaptic connection with ASH and the command interneuron AVA, and also with sleep-induction neuron RIS, also showed no sleep-dependent difference in response. Together, these results suggest that modulation in sensory processing that leads to reduced arousal response in sleep occurs presynaptic to AVA, either from ASH, AIB, or another interneuron (**Fig. S5c**), or perhaps via neuropeptides from other sources. One possibility is that sleep increases arousal threshold predominately by diminishing the efficacy of monosynaptic shortcuts to the command interneurons (here, ASH to AVA), whereas sensory information is preserved to first layer interneurons (such as AIB) to allow for rapid arousal from more salient polymodal stimuli from multiple sensory neurons. However, animal survival should benefit from maintaining rapid arousal to potentially harmful stimuli, yet for aversive ASH sensory neurons, this does not appear true. Alternatively, the dampened brain state apparent in sleep^74^ may broadly suppress activity in premotor interneurons like AVA, increasing arousal thresholds equally to all types of sensory input. These data highlight the hierarchy of altered sensorimotor processing, and further study of response modulation to additional sensory stimuli should provide insight into the architecture and mechanisms of sleep-dependent modulation of arousal.

These flexible microfluidic systems for studying adult sleep in *C. elegans* are applicable to any neuron, stimulus, environment, and genetic perturbation for thorough assessment of sleep behavior and underlying neural responses. For example, it will be informative to compare neural responses in various sleep modes, including hypoxia and starvation-induced sleep as shown here, as well as heat shock and satiety-related sleep. Microfluidic devices are easily customized to different animal sizes by adjusting arena post geometry, for example, to observe L4 animals in lethargus transition stages in developmentally-timed sleep. Other types of oxidative or metabolic stress (such as by chemical oxidants or varying food quality), or sleep disruption via mechanical stimulation or light, can be applied using the same microfluidic devices and tracking methods. Overall, this platform can be used to uncover molecular and neural circuit pathways underlying altered sensation during sleep, toward establishing connections between nematode sleep and associated regulatory mechanisms and human sleep disorders.

## Acknowledgements

We thank R. Lagoy, K. Burnett, H. White, L. Innarelli, E. Larsen, A. Marley, J. Srinivasan, and D. Colón-Ramos for experimental support and feedback. We thank J. Florman and M. Alkema for graciously providing the AVA∷GCaMP imaging line. Some strains were provided by the CGC, which is funded by NIH Office of Research Infrastructure Programs (P40 OD010440). D.R.A. was supported by NSF CBET 1605679 and EF 1724026, NIH R01DC016058, and a Career Award at the Scientific Interface (CASI) from the Burroughs Wellcome Fund. D.E.L. was supported by an NSF IGERT award (DGE 1144804). Y.L.C. was funded by an EMBO Long-term Fellowship (ALTF 403-2016). Research in the D.A.C-R. lab for A.A. and J.D.H. was supported by a Whitman Fellowship from the Marine Biological Laboratories, NIH R01NS076558, DP1NS111778 and by an HHMI Scholar Award. J.D.H. was supported by the Ruth L. Kirschstein NRSA (NIH F32MH105063).

## Author contributions

D.E.L. designed and performed experiments, developed methods, analyzed and interpreted data, and wrote the paper. Y.L.C., J.D.H. and W.R.S. interpreted data and developed reagents. A.A. developed reagents. D.R.A. designed experiments, developed methods, analyzed and interpreted data, and wrote the paper.

## Ethics declarations

The authors declare no competing interests.

## Methods

### Strains and *C. elegans* culture

All *C. elegans* strains were maintained under standard conditions on NGM plates and fed OP50 *E. coli* bacteria seeded onto each plate. Wild-type animals were Bristol strain (N2). The following mutant strains were used: CB1611, *mec-4 (e1611)*; FK103, *tax-4 (ks28)*; CX32, *odr-10 (ky32)*. Neural imaging strains expressing GCaMP in specific neurons were: (ASH^52^) CX10979, *kyEx2865* [*Psra-6∷GCaMP3; Pofm-1p∷GFP*]; (AIB) DCR6035, *olaIs94* [*Pinx-1∷GCaMP6f; Punc-122∷GFP*]; (AVA) QW607, *zfls42* [*Prig-3∷GCaMP3∷SL2∷mCherry*] gifted by the Alkema lab; (RIS) AQ4064, *ljEx1119* [*Pflp-11∷GCaMP3∷SL2-tagRFP;unc-122∷rfp*]. To make the RIS imaging line, a 2643 bp region immediately upstream of the ATG of the *flp-11* gene was amplified, similar to previously reported methods^73^. This promoter was shown to express consistently in RIS and occasionally in other neurons^73^. To synchronize for age, we picked L4 larval stage animals one day prior to experimentation such that all animals tested were at the young adult stage.

To prepare for an experiment, animals were isolated by genotype and/or arena placement and then transferred to an unseeded NGM plate immediately prior to experimentation. The plates were then flooded with the control buffer used for their respective experiment: S. Basal buffer (100 mM NaCl, 50 mM KPO_4_; pH 6.0) for unfed behavioral experiments, S. Medium buffer (1 L S. Basal, 10 mL 1 M potassium citrate pH 6.0, 10 ml trace metals solution, 3 ml 1 M CaCl_2_, 3 ml 1 M MgSO_4_) for feeding experiments, or a saline buffer (80 mM NaCl, 5 mM KCl, 20 mM D-glucose, 10 mM HEPES, 5 mM MgCl_2_, 1 mM CaCl_2_; pH 7.2) for copper chloride stimulus experiments. Animals were then collected into loading tubing using a 1 mL syringe prior to injection into the microfluidic arena.

### Microfluidic device fabrication

“Population behavior” and “Neural imaging” microfluidic devices were fabricated as previously described^74^. Briefly, transparency photomasks were printed at 25,000 dpi from designs sketched using DraftSight CAD software. SU-8 mold masters were prepared on silicon wafers using standard photolithography techniques, and microfluidic devices were fabricated by pouring degassed PDMS (Sylgard 184, Dow Corning) onto the mold and heat curing. Individual devices were then cut out and punched to provide inlet and outlet flow. A hydrophobic glass substrate was created by vapor deposition of tridecafluoro-1,1,2,2-tetrahydrooctyl trichlorosilane (TFOCS, Gelest) and then sealed reversibly to the microfluidic channels. An upper glass slide, with holes drilled over inlet and outlet ports with a diamond-coated drill bit, was sealed above the device, which was then was placed into a metal clamp.

### Stimulus preparation

All odor dilutions were freshly prepared on the day of experimentation. NA22 *E. coli* stock solutions were prepared using previously described methods^75^. Briefly, NA22 *E. coli* was cultured, concentrated into pellet form, and suspended in S. medium buffer. A stock solution was diluted to an OD600 of 7.0, and 50 μg/ml of kanamycin was added to prevent bacteria from growing. Chemical solutions were prepared at a 1:20 dilution of stock solution and filtered through a 5 μm filter. Diacetyl (1.1 μM) was prepared from a 10^−3^ dilution (11 mM) stock solution immediately prior to experimentation. Serotonin was prepared by dissolving serotonin creatine sulfate monohydrate powder (Sigma). Sodium sulfite (Sigma) solution was prepared moments before experimentation at 30 mM. We found that a 30 mM sodium sulfite solution would remain at nearly 0% oxygen with stirring for 12 h and without stirring for 5 days (Ocean Optics Neofox O_2_ probe kit), so the testing solution would be devoid of oxygen for entire 12 h testing period. The control solution of sodium sulfate was created by allowing for reoxygenation of the sodium sulfite solution for greater than 5 days. For neural imaging experiments, 1 mM copper chloride solution was prepared the day of the experiment using copper chloride powder.

### Microfluidic device setup

Microfluidic devices were cleaned, assembled, and degassed in a vacuum desiccator for 30–60 min prior to experimentation. Degassing devices accelerates the absorption of air bubbles within the device. For behavioral experiments, devices were filled with 5% (w/v) Pluronic F127 through the outlet port to prevent bacterial and molecular absorption by passivation of the microfluidic surfaces and to minimize bubble entrapment via its surfactant properties. Neural imaging devices were filled with control buffer alone. Reservoirs of loading solutions were prepared as previously described^74^, purging the reservoir system of bubbles and connecting the tubing into the inlets of the device. Once flow was properly established, animals were gently loaded into their respective arenas and allowed to roam for 15–20 min prior to experimentation. For neural imaging experiments, a control valve is used to switch between stimulus and control buffer conditions within 0.5 s (**Fig. S4d**).

### Population behavior imaging and identification of sleep events

Videos of population behavior were captured using a 6.6 MP PixelLink FireWire Camera at 1 fps for 12 h with an image resolution of ~30 pixels/mm. Videos were processed after experimentation as previously described using MATLAB to extract behavioral data^51^, and then further analyzed to identify sleep events. A minimum sleep entry window of 20 s and exit window of 5 s were used to quantify state transitions. To verify accuracy in parameters for sleep detection, user observed behavioral state was compared to script calculated state on randomly chosen 60 second traces of an individual animals (**Fig. S1e**). All behavior data was collected using “Population behavior” devices with four 16 mm × 15 mm arenas capable of housing ~25 animals per arena for simultaneous study.

### Neural calcium imaging, sleep detection and data analysis in closed-loop system

Closed-loop neural imaging videos were acquired at 5× magnification (NA=0.25) with a Hamamatsu Orca-Flash 4.0 sCMOS camera using MicroManager/ImageJ software. The system has a green (λ = 520-550 nm) LED mounted overhead to provide pulsed brightfield illumination for tracking animal behavior and a Lumencor SOLA-LE solid-state lamp pulsed to excite GCaMP during fluorescence calcium imaging. To achieve autonomous experimentation for a closed-loop system, custom Arduino, MicroManager, and ImageJ scripts work together to control illumination timing, image acquisition, stimulus delivery, and sleep/wake state identification. An Arduino Uno microcontroller was programmed to control fluidic valves through a ValveLink 8.2 (AutoMate Scientific) controller and to control illumination sources for brightfield and fluorescent imaging. A MicroManager script allows the user to configure all camera and illumination settings prior to experimentation as well as all testing conditions for sleep assessment. Once the experiment is underway, the script initiates brightfield image capture at the desired framerate, and analyzes movement compared with the prior image in real time to determine the animal’s behavioral state. If the current state and timing match the desired and preprogrammed conditions for neural imaging, the script initiates a fluorescence image stack recording and communicates with the Arduino via serial commands to control epifluorescence illumination and chemical stimulation with the desired timing.

Tracking of behavior of a single animal in the closed-loop neural imaging system was done using brightfield illumination with images captured at 0.1 fps. The current sleep/awake state of the animal was determined by an ImageJ script which calculates a movement index for each frame, represented as the fraction of body pixels moved since the previous frame, ranging from 0 – 1 (**Fig. S4a**). A sleep state was defined as movement below the empirically-optimized threshold (0.125) for 3 consecutive frames (i.e., for 20–30 s). Optimization of detection parameters was done by maximizing accuracy from user observed behavioral states to script calculated states (**Fig. S1e**). One minute of consistent sleep or wake state frames were used to increase confidence in the animals’ current state before neural imaging.

Calcium imaging was performed on freely-moving animals as previously described^52^ using lines expressing GCaMP in selected neurons. Neural activity was recorded in RIS neurons at 2 fps with no stimulation from the closed-loop system, however motion was detected post-processing to identify sleep bouts. Calcium imaging in ASH, AIB, and AVA neurons was performed at 10 fps, using closed-loop stimulation to record responses to 10 s chemical stimulation from 5–15 s within a 30 s trial. Videos were analyzed for neural fluorescence and locomotion using NeuroTracker software in ImageJ, which tracks the position of the neuron over time and integrates fluorescent intensity of the soma using a 4 × 4 pixel box. Fluorescence (*F*) was normalized by dividing by the initial baseline fluorescence in the first 4 s of each trial before stimulation (*F*_0_). As AIB fluorescence may not be at baseline at the beginning of each trial, baseline AIB intensity was determined for each animal across all trials, and individual AIB traces were excluded when animals engaged in reversal behavior immediately prior to stimulation.

### Statistical analysis

Statistics were performed using one-way ANOVA with Bonferroni’s correction for multiple comparisons or an unpaired two-tailed t-test when specified for 2 sample comparison, using the Statistics and Machine Learning Toolbox in MATLAB. Data represented as mean ± s.e.m. unless otherwise stated. In behavioral experiments, animals were excluded when valid behavioral tracks comprised <8% of recording time, indicating an animal not viable or not present during the test. In neural recordings, the top and bottom 1% of instantaneous fluorescent intensity was removed to reduce noise in peak fluorescence calculations.

### Data Availability

The data that support the findings of this study are available from the corresponding author upon reasonable request.

### Code Availability

Scripts and code for control systems and data analysis are available upon request.

## Supplementary Information

**Supplementary Figure 1.**
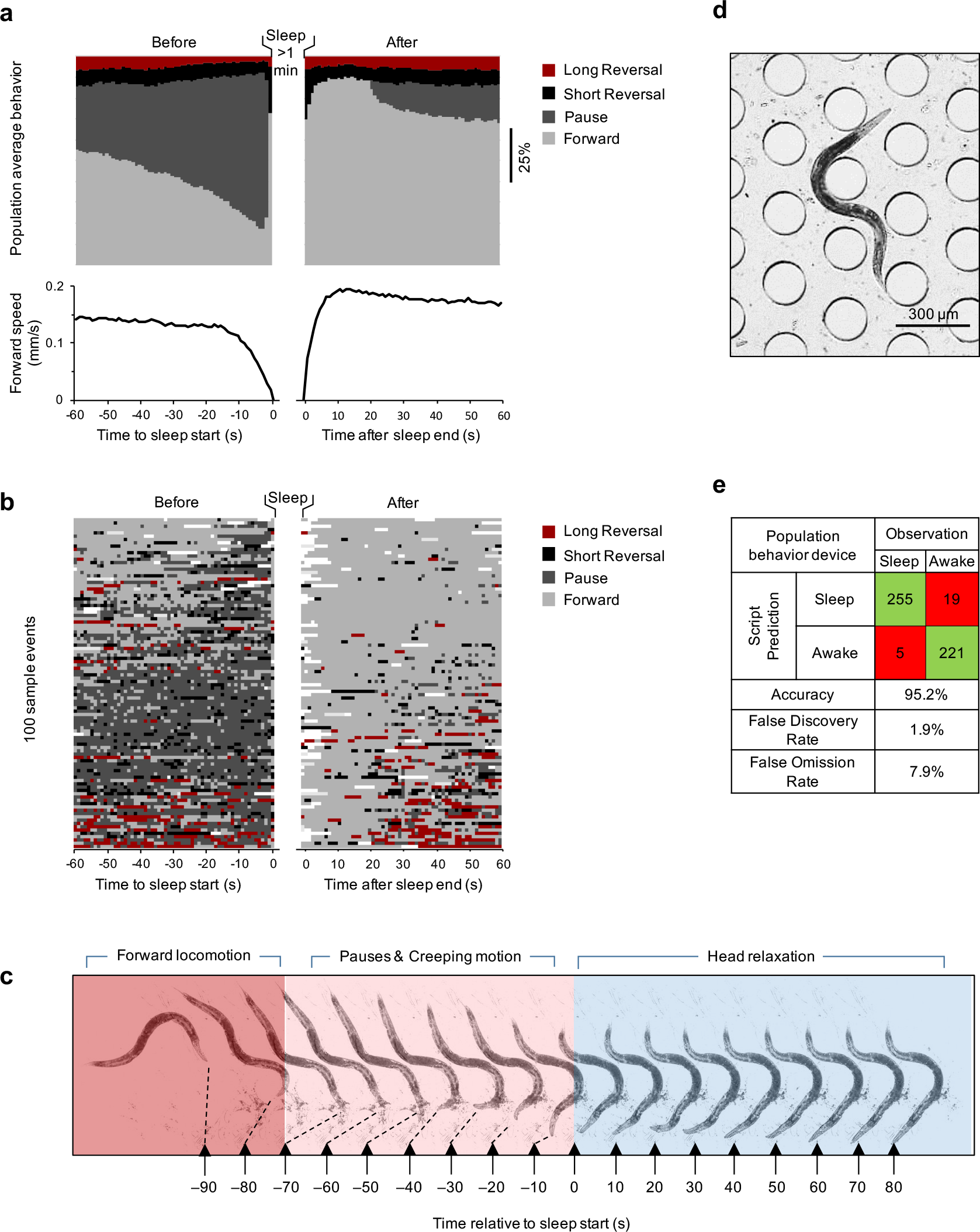
Spontaneous adult sleep transitions in microfluidic devices. (**a**) Distribution of behavior probability and average speed in the 60 s before and after a sleep bout of at least 1 minute. Data are from 697 wild-type (N2) animals observed for 12 h (4359 qualified bouts, 32% of all sleep bouts >60 s). Examples exclude sleep bouts that preceded or followed another sleep bout within 60 s. (**b**) Ethogram of behavior state of 100 randomly selected animals in the 60 s before and after a sleep bout, sorted such that animals with more frequent forward behavior pre-sleep are at the top, and with more prevalent turning behavior at the bottom. White coloring denotes unknown behavior. (**c**) Montage of an animal transitioning between awake forward motion (red), pausing/creeping motion (pink) and sleep with characteristic head relaxation (blue). (**d**) Image of a sleeping animal in the microfluidic device. Head exhibits straight posture, as opposed to curling around posts as seen in awake animals. (**e**) Table of accuracy verification for automatic sleep tracking in the “Population behavior device”.

**Supplementary Figure 2.**
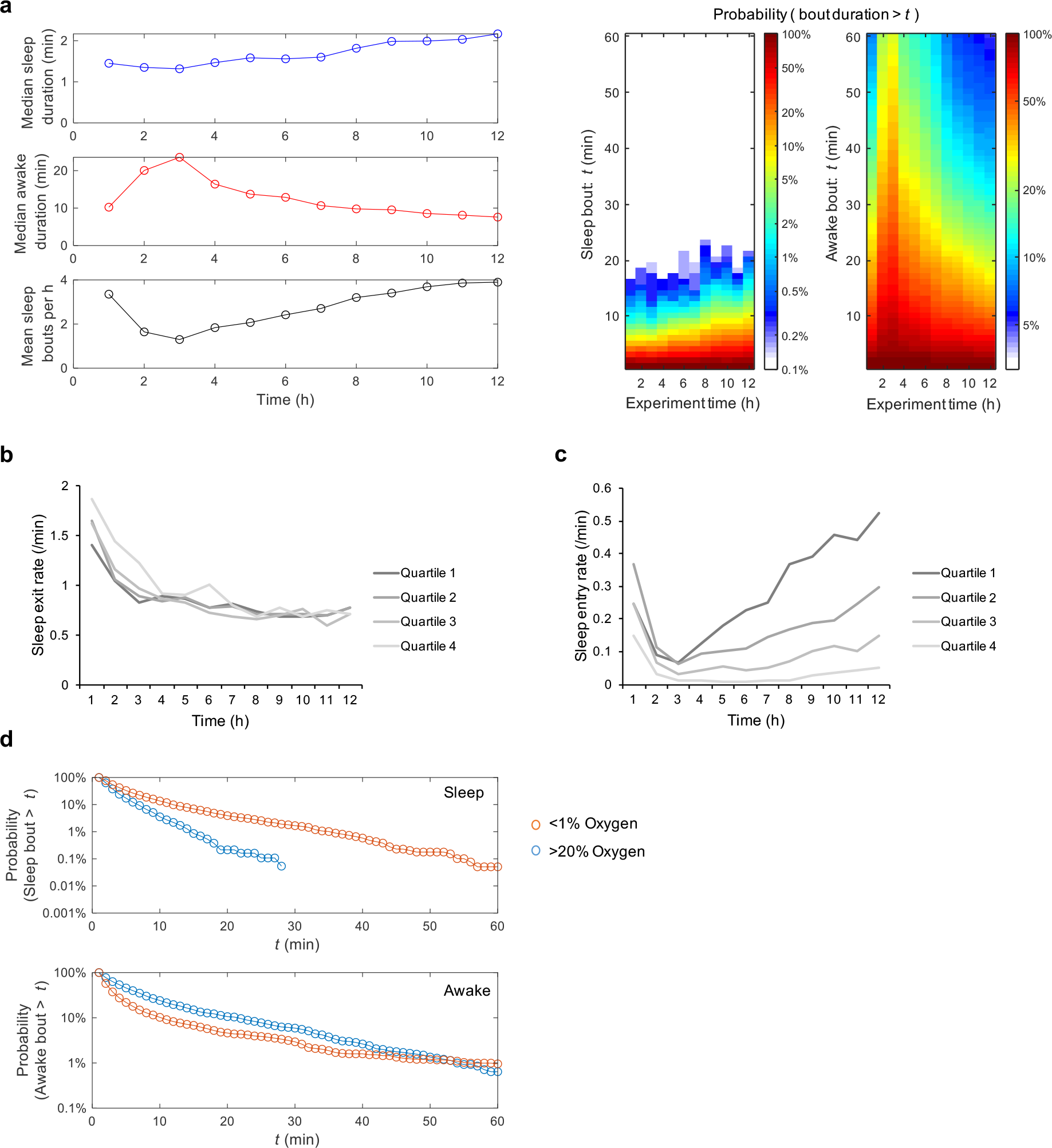
Characteristics of spontaneous wild-type adult sleep and wake periods. (**a**) Dynamics of sleep behavior over 12 h from Fig. 1c. Median sleep bout duration, median awake bout duration, and average number of sleep bouts plotted per hour. Heatmap shows probability (%) of sleep and awake bout durations greater than the specified times. (**b**) Sleep exit rate (average transitions per minute) as in Fig. 1f for each total sleep quartile in Fig. 1d,e, showing little inter-quartile variation from most sleep (quartile 1) to least sleep (quartile 4). (**c**) Sleep entry rate for each total sleep quartile in Fig. 1d,e, showing large inter-quartile variation across wild-type animals. (**d**) Sleep and awake bout duration distributions for high and low oxygen from data in Fig. 2a, plotted as probability of durations larger than the specified times. Low oxygen increases sleep duration and frequency, and shortens awake periods.

**Supplementary Figure 3.**
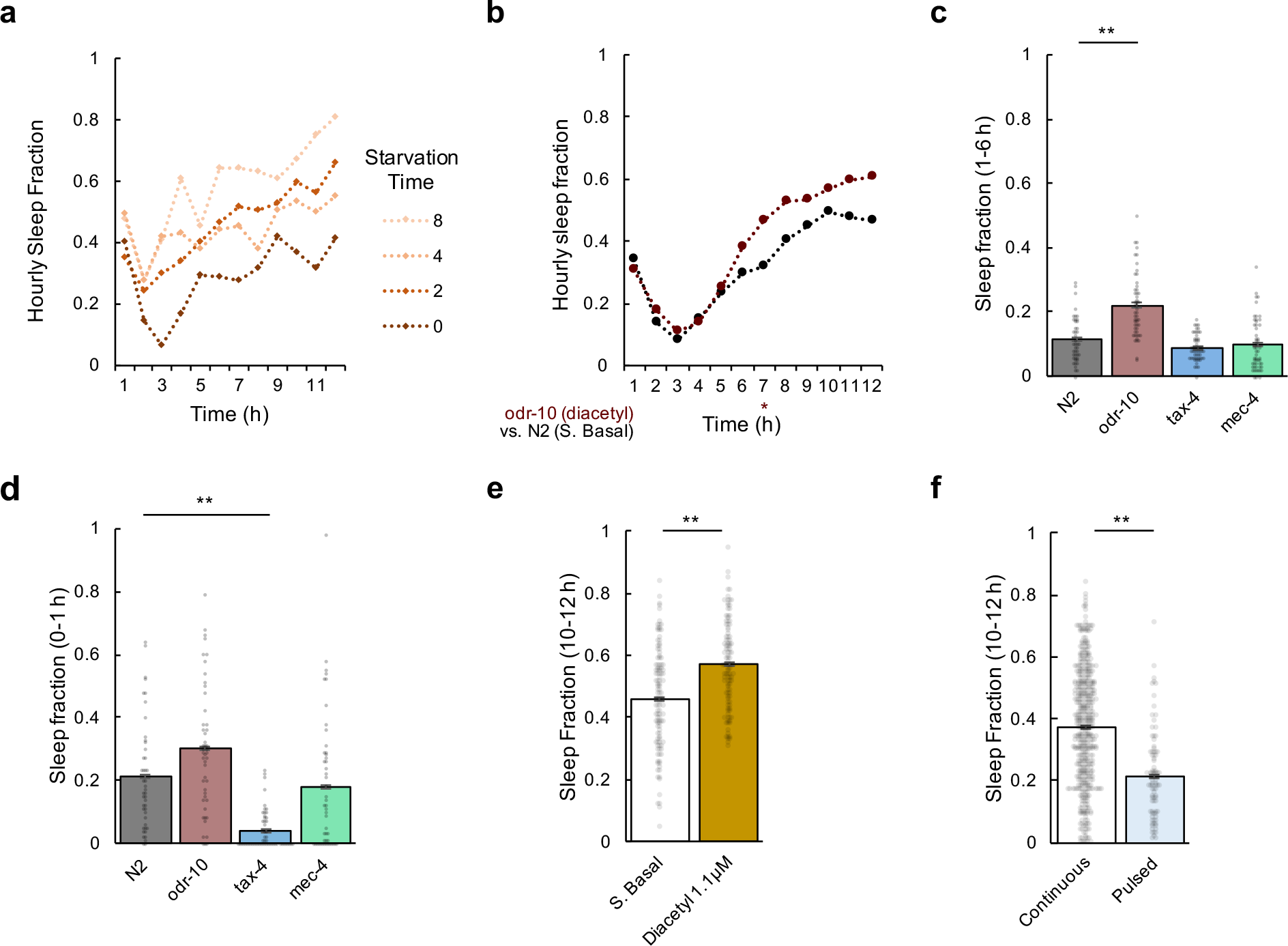
Comparisons of sleep fraction among specific groups and time ranges. (**a**) Hourly sleep fraction comparison following wild-type starvation times from 0–8 h on agar dishes prior to loading in the microfluidic device. (**b**) Hourly sleep fraction comparison between *odr-10* mutants under constant exposure to 1.1 μM diacetyl from Fig. 2g to N2 animals in S. Basal from Fig. 2d. (**c**) Total sleep fraction from hour 1 to hour 6 from data in Fig. 2g. (**d**) Total sleep fraction during the first hour of experiment from data in Fig. 2g. (**e**) Total sleep fraction during hours 10–12 comparing S. Basal and diacetyl conditions from data in Fig. 2d. (**f**) Total sleep fraction during hours 10–12 comparing continuous flow of S. Basal (Fig. 1c) vs. pulsed flow while flow is on (Fig. 1h). Statistics for **b** were performed using one-way ANOVA with Bonferroni’s correction for multiple comparisons. For hourly sleep fraction, significance is noted for * P<0.0001 as indicated within data of that hour. Statistics for plots **c** and **d** were performed using one-way ANOVA with Bonferroni’s correction for multiple comparisons; ** P<0.001. Statistics for plots **e** and **f** were performed using an unpaired two tailed t-test; ** P<0.001.

**Supplementary Figure 4.**
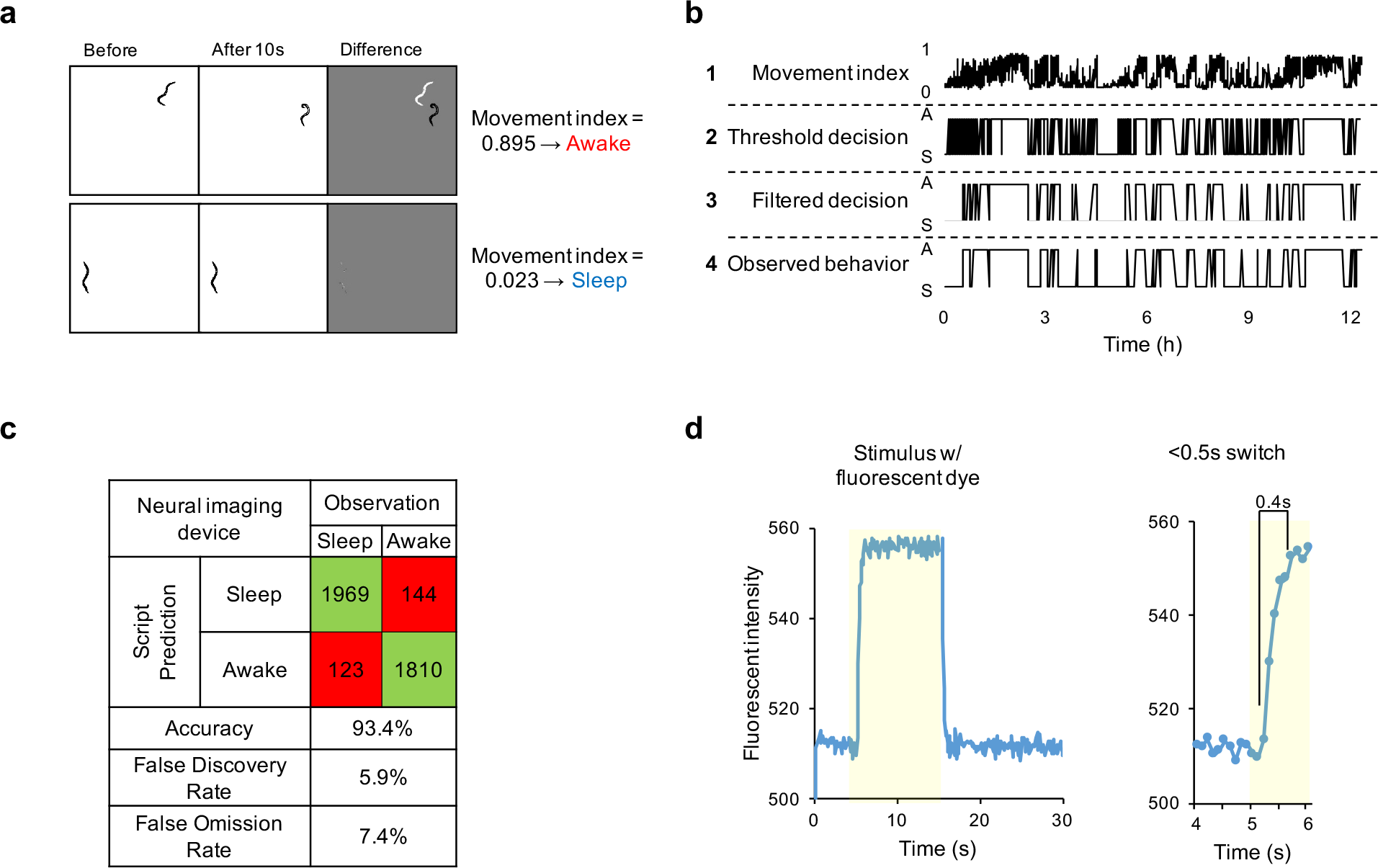
Sleep detection in “Neural imaging” microfluidic devices. (**a**) Examples of the frame subtraction method for sleep detection showing awake and sleep cases. Movement Index represents fraction of body pixels moved between 10 s frames. (**b**) Schematic of sleep decision processing in a single wild-type animal over 12 h in S. Basal, top-to-bottom: 1. Movement Index (MI); 2. Result of threshold MI < 0.125; 3. Temporal-filtering for 5 consistent state frames (40 s); 4. Human observation. (**c**) Table of accuracy verification for automatic sleep tracking in the “Neural imaging” device. (**d**) Raw fluorescent pixel intensity measurement over time from a 5–15 s pulse of fluorescent dye, switching within 0.4 s (right).

**Supplementary Figure 5.**
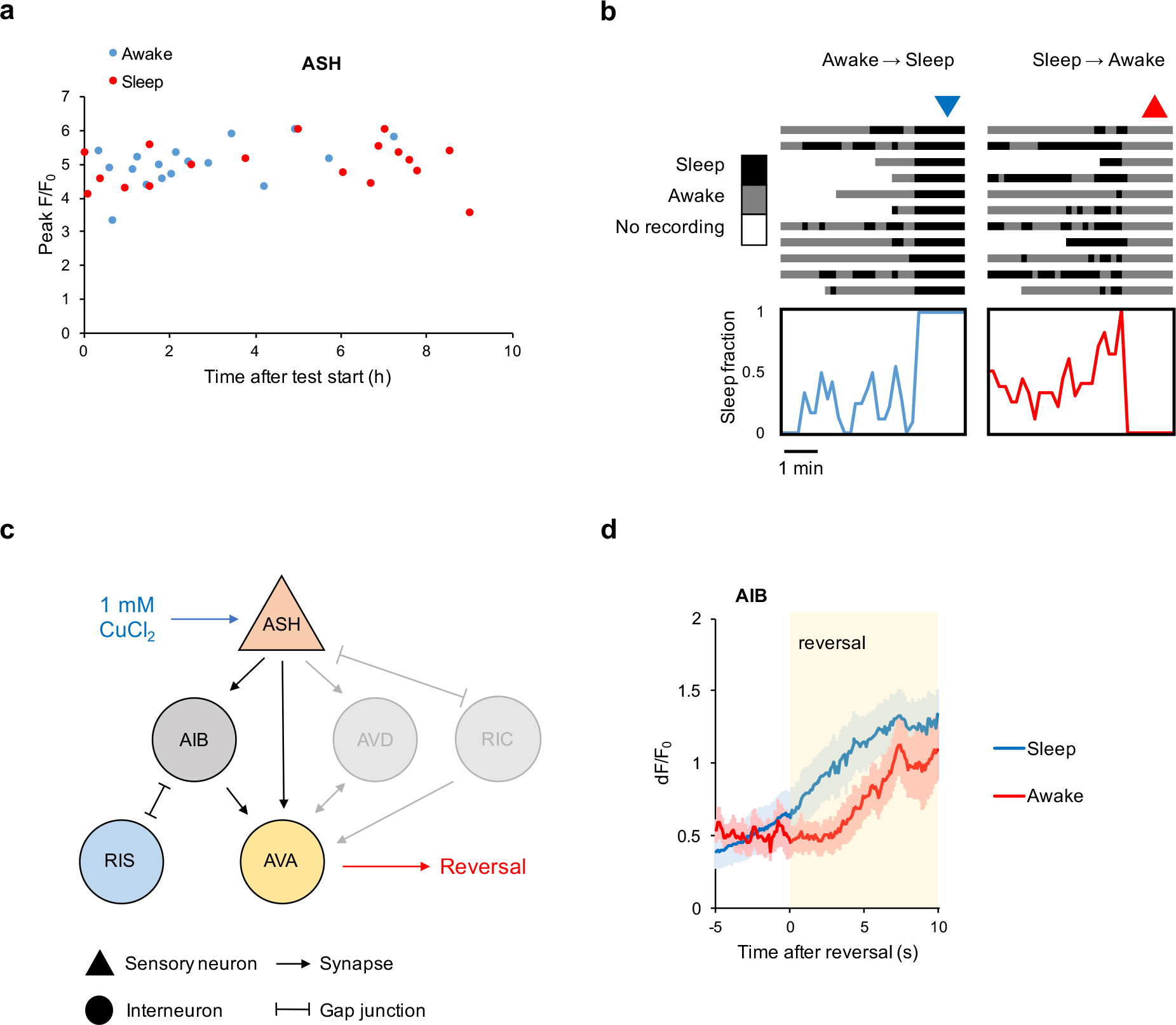
Neural imaging during closed-loop experiments. (**a**) Peak F/F_0_ response in ASH to 1 mM CuCl_2_ plotted by time after animal started closed loop experiment. Data from Fig. 4e,f. (**b**) Pre-stimulus behavior up to 5 min before 22 stimulation trias during an example closed-loop experiment from Fig. 4d. Average sleep fraction is plotted below. (**c**) Neural connections linking copper chloride sensation for reversal behavior upon arousal by stimulus in sleep and wake states in Fig. 4e-j. (**d**) Neural activity from AIB aligned to point of first reversal response. Data from Fig. 4g,h. Lines indicate mean response; shading s.e.m.

### Supplementary Video Legends

**Supplementary Video 1. Sleep tracking of wild-type adult** *C. elegans* **in a “Population behavior” device.** Video shows three time periods of a 12 h video tracking sleep behavior in wild-type animals: early (~1.5–2 h), mid (~5.5–6 h), late (11–11.5 h). Animals automatically detected in a sleep state are indicated with a blue circle. Video is accelerated 100x, with time stamp (h:m:s) in upper left corner. Video frame is 32 mm × 33 mm (w × h).

**Supplementary Video 2. Example of closed-loop, sleep state-triggered chemical stimulation and ASH sensory neuron recording in a “Neural imaging” device.** Video shows an animal expressing GCaMP in the ASH neuron tracked in the closed loop system every 10 s to determine sleep state, then stimulated with a 1 mM copper chloride pulse (10 s duration) 1 min after each sleep state change. First, an awake animal is detected and a stimulus paradigm is applied, with fluorescence recording at 10 fps. Next, a sleep bout is detected and ASH neural response recorded. A timestamp (h:m:s) is shown in upper left and state change detections and CuCl_2_ pulses are indicated. Video is accelerated 37.5× during brightfield state determination and 3× during neural response recording. Video frame is 3.6 mm × 2.7 mm (w × h).

